# Guided Visual Search is associated with a feature-based priority map in early visual cortex

**DOI:** 10.1101/2024.10.03.616568

**Authors:** Katharina Duecker, Kimron L. Shapiro, Simon Hanslmayr, Benjamin J. Griffiths, Yali Pan, Jeremy Wolfe, Ole Jensen

**Affiliations:** Centre for Human Brain Health, School of Psychology, University of Birmingham, UK; Department of Neuroscience, Brown University, RI, USA; Centre for Cognitive Neuroimaging, School of Neuroscience and Psychology, University of Glasgow, UK; Brigham and Women’s Hospital, Boston, MA, USA; Harvard Medical School, Boston, MA, USA; Department of Experimental Psychology, University of Oxford, UK; Oxford Centre for Human Brain Activity, Wellcome Centre for Integrative Neuroimaging, Department of Psychiatry, University of Oxford, UK

## Abstract

Visual search models have long emphasised that task-relevant items must be prioritised for optimal performance. While it is known that search efficiency also benefits from active distractor inhibition, the underlying neuronal mechanisms are debated. Here, we used MEG in combination with Rapid Invisible Frequency Tagging (RIFT) to probe how neuronal excitability in early visual cortex is modulated during feature-guided visual search. Participants were instructed to indicate the presence or absence of a letter “T” presented amongst 16 and 32 “L”s. In the *guided search* condition, participants were informed about the colour of the “T” and could infer the colour of the irrelevant distractors. In the *unguided search* condition, the target colour was unknown. We found that *guided search* was associated with enhanced RIFT responses to the target colour, and decreased responses to the distractor colour compared to *unguided search*. These results conceptually replicated using both a conventional coherence approach, as well as with a General Linear Model approach based on a single-trial measure of the RIFT response. The present findings expand on previous reports based on electrophysiology and fMRI in humans and non-human primates by revealing that feature-guidance in visual search affects neuronal excitability as early as primary visual cortex.

## Introduction

Visual search is a widely used paradigm, applied to operationalise the everyday task of finding a pre-defined stimulus (target) among distracting stimuli (distractors), for instance, a friend in a crowd. Search is more efficient when low-level features of the target, e.g., colour or shape, are known to the observer. For example, when we know that our friend is wearing a yellow raincoat, we will pay less attention to people wearing blue jackets. The allocation of visual attention has long been suggested to involve a priority map: a representation of objects in the visual field in which object locations are weighted based on their saliency and task-relevance (Awh et al., 2012; Koch & Ullman, 1985; Navalpakkam & Itti, 2005; Serences & Yantis, 2006; Thompson & Bichot, 2005; Zelinsky & Bisley, 2015). Priority maps have become a central component of models of selective attention (Awh et al., 2012; Bisley, 2011; Fecteau & Munoz, 2006) and visual search (Wolfe, 1994, 2021). In the example above, this map would assign high priority to objects containing the colour yellow and low priority to those containing the colour blue.

Evidence from behavioural, electrophysiological, and neuroimaging studies leaves little doubt that visual attention is guided by a mechanism akin to a priority map, whereby neural responses to the target are boosted, and responses to the distractors are reduced or suppressed (Andersen et al., 2008; Bayguinov et al., 2015; Bichot & Schall, 1999; Bisley & Goldberg, 2010; Bisley & Mirpour, 2019; Chelazzi et al., 1993, 2014; Cosman et al., 2018; Fecteau & Munoz, 2006; Gottlieb et al., 1998; Hickey et al., 2009; Ipata, Gee, Goldberg, et al., 2006; Ipata et al., 2009; Klink et al., 2023; Li, 2019; Luck & Hillyard, 1994b, 1994a, p. 2006; Mirpour et al., 2009; Motter, 1994; Müller et al., 2006; Ptak, 2012; Serences & Yantis, 2006; Sprague & Serences, 2013; Thompson et al., 2005; Thompson & Bichot, 2005). Modulation of cortical excitability in accordance with a priority map has, for instance, been observed in electrophysiological recordings in non-human primates from the frontal eye field and lateral intraparietal cortex (Cosman et al., 2018; Ipata, Gee, Goldberg, et al., 2006; Ipata, Gee, Gottlieb, et al., 2006) as well as V4 (Klink et al., 2023). Traditionally, neuronal excitability in primary visual cortex has been argued to merely underlie the saliency of the sensory input, i.e. implementing a saliency map (Itti & Koch, 2001; Li, 2002; Melloni et al., 2012). However, recent studies have provided evidence for the activity in early visual areas being modulated by attention; as e.g. quantified by the BOLD signal in fMRI (Beffara et al., 2023; Foster & Ling, 2022; D. Richter et al., 2024) and intracranial recordings in non-human primates (Yin et al., s). These findings align with the notion that involving V1 in guiding the search be beneficial due to a high spatial resolution of foveal vision (Hochstein & Ahissar, 2002; Li, 2019). Here, we use MEG and rapid photic stimulation in a classic guided search paradigm to test this idea with high temporal precision.

Behavioural studies of visual search often involve complex search displays with a large number of stimuli (Wolfe, 2020, 2023). Electrophysiological approaches in humans and non-human primates, however, rely on spatially separable stimuli and therefore typically investigate visual search paradigms with smaller set sizes of up to six items (Donohue et al., 2018; Forschack et al., 2022; Hickey et al., 2009; Ipata et al., 2009; Klink et al., 2023; van Zoest et al., 2021). Other studies extrapolate the underlying mechanisms of visual search from experiments on selective attention, cueing the participant to attend to certain objects presented in a large field of stimuli (Andersen et al., 2008; Andersen & Müller, 2010; Müller et al., 2006). The neural dynamics of distractor suppression in humans and non-human primates are typically investigated in the context of actively ignoring a single, salient distractor (Donohue et al., 2020; Feldmann-Wüstefeld et al., 2020, 2021; Ferrante et al., 2023; Forschack et al., 2022; Gaspar & McDonald, 2014; Gaspelin & Luck, 2018; Hickey et al., 2009; Jannati et al., 2013, 2013, 2013; Sawaki & Luck, 2010; Serences et al., 2004; van Zoest et al., 2021). In the context of these studies, it has long been debated whether the location of an expected singleton distractor can be suppressed in anticipation of the search display (Luck et al., 2021), with several studies arguing both for (D. Richter et al., 2024) and against (Klink et al., 2023) anticipatory distractor suppression in visual cortex.

Here, we leveraged MEG in combination with Rapid Invisible Frequency Tagging (RIFT) to investigate target boosting and distractor suppression in a classic visual search paradigm. In the tradition of early behavioural studies that motivated the hypotheses that search is guided by a map of the visual field with 16 and 32 coloured stimuli (Treisman et al., 1980; Wolfe, 1994, 2021) (Figure 1a). RIFT is a novel, subliminal stimulation method to probe neuronal excitability in early visual cortex, while leaving endogenous spontaneous oscillations unperturbed (Duecker et al., 2021; Spaak et al., 2024; Zhigalov et al., 2019; Zhigalov & Jensen, 2020). The high spatial and temporal resolution of the MEG recording, paired with the high frequency range used for RIFT, allowed us to estimate both the source of the RIFT signal, and the latency of the attention effects. As we will show, the RIFT responses demonstrate that both target boosting and distractor suppression affect neuronal excitability as early as V1 (also see Bouwkamp et al., 2023). Based on the time course of the RIFT signal, we conclude that this modulation underlies downstream control from higher-order visual areas, such as the frontal eye field, lateral intraparietal cortex, and V4 (Cosman et al., 2018; Ipata, Gee, Goldberg, et al., 2006; Ipata, Gee, Gottlieb, et al., 2006; Klink et al., 2023).

**Figure 1.**
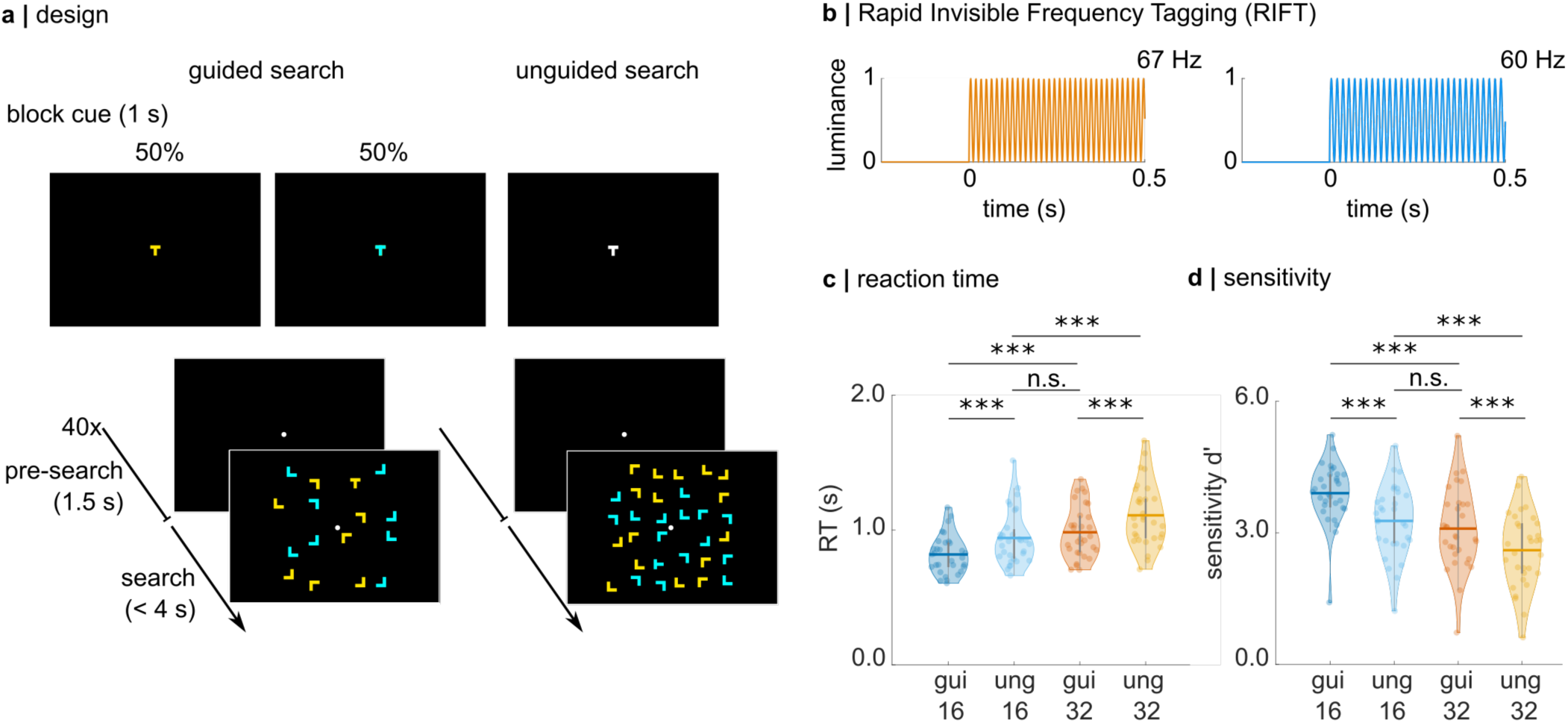
Experimental paradigm, Rapid Invisible Frequency Tagging (RIFT), and search performance. **a** Trials were presented in a blocked design. Each block contained 20 target absent and 20 target present trials. Set sizes (16 or 32) were the same within each block. At the start of a block in the guided search condition, a “T” was presented in yellow or cyan, revealing the target colour for the following 40 trials. A block in the unguided search condition began with the presentation of a white “T”, and the target colour was randomized over trials. Note that the search displays are not true to scale; the eccentricity of the search array amounted to 10° visual angle, 5° on either side of the fixation dot. **b** RIFT at 60 and 67 Hz was applied to the colour of the stimuli by modulating the luminance sinusoidally. In this example, yellow stimuli were tagged at 67 Hz and cyan stimuli were tagged at 60 Hz. **c** Search performance decreases for more difficult searches. A hierarchical regression approach reveals a significant main effect for set size (β = 0.180) and guided/unguided (β = - 0.138). Indicating that larger set sizes are associated with slower responses, while guided searches are faster than unguided searches. Pairwise comparisons reveal no significant difference in reaction time between unguided search set size 16 and guided search set size 32 (V = 134, z = 2.24, r=0.4, p = 0.148), suggesting that participants focused their search on task-relevant items in the guided search condition. **d** Analogously, for accuracy (as measured by d’), hierarchical regression reveals a significant main effect for set size (β = - 0.74) and guided/unguided (β = 0.56), indicating that accuracy is higher in guided searches and for set size 16 compared to 32. Again, there is no significant difference in sensitivity for unguided search set size 16 and guided search set size 32 (t(30) = 2.2, d= 0.2, p = 0.23).

## Results

Our experimental paradigm featured two search conditions (*guided* and *unguided search*) and two set sizes (16 and 32), presented in a block design (with a randomised order over all participants), with each block consisting of 40 trials (Figure 1a). Participants were instructed to indicate if a single letter “T” was presented among several “L”s. In the *guided search* condition, participants were cued to the colour of the target “T” (either yellow or cyan) at the beginning of the block. Importantly, as only these two colours were used throughout the experiment, participants were able to infer the distractor colour from this cue. In the *unguided search* condition, a white “T” was presented at the beginning of the block, meaning the target and distractor colours were not cued, and the colour of the T was randomised over trials. Set size was kept constant within each block. The target and distractor colours were frequency-tagged by modulating their luminance sinusoidally, at 67Hz and 60Hz respectively (balanced over trials; Figure 1b). Participants were instructed to perform the task while fixating on a centrally presented dot. Note that the 60 and 67 Hz flicker are invisible to the observer but modulate neuronal activity (Spaak et al., 2024).

Based on the extensive literature on visual search, we predicted search performance to be worse (indicated by reaction time and accuracy *d’*) for more difficult searches, i.e., *set size 32* relative to *16* and *unguided* compared to *guided search* (Egeth et al., 1984; J. Palmer, 1994; Wolfe, 1994, 2021). Indeed, these hypotheses were confirmed by a hierarchical regression approach applied to the average reaction time and sensitivity (d’) for each participant. Reaction time was significantly increased for higher set size (β = 0.180) and was reduced for *guided* compared to *unguided search* (β = - 0.138, see Supplementary analyses). A Wilcoxon-signed rank test revealed no significant difference between *unguided search, set size 16* and *guided search, set size 32*, indicating that the difficulty of these searches was similar as would be expected if colour guidance could render half of the distractors irrelevant in the *guided search, set size 32* condition (V = 134, z = 2.24, r=0.4, p = 0.148, see Supplementary Table 2, Supplementary Table 4, and Figure 1c). Similarly, sensitivity decreased for the higher set size (β = - 0.74) and increased for guided vs unguided search (β = 0.56), with no difference between *unguided search, set size 16* and *guided search, set size 32 as indicated by an independent sample t-test* (t(30) = 2.2, d= 0.2, p = 0.23). These behavioural findings demonstrate that the participants used the colour cue at the beginning of the block to focus their search on the target colour.

### Rapid Invisible Frequency Tagging responses indicate target boosting and distractor suppression

RIFT elicited brain responses at the respective stimulation frequencies that were detected in a small number of MEG sensors over the occipital cortex (Figure 2a, left, see Supplementary Figure 1 for individual topographic representations per participant). Source modelling based on Dynamic Imaging of Coherent Sources (DICS) demonstrated that the responses emerged from early visual regions (V1, MNI coordinates [0 -92 -4], Figure 2b).

**Figure 2.**
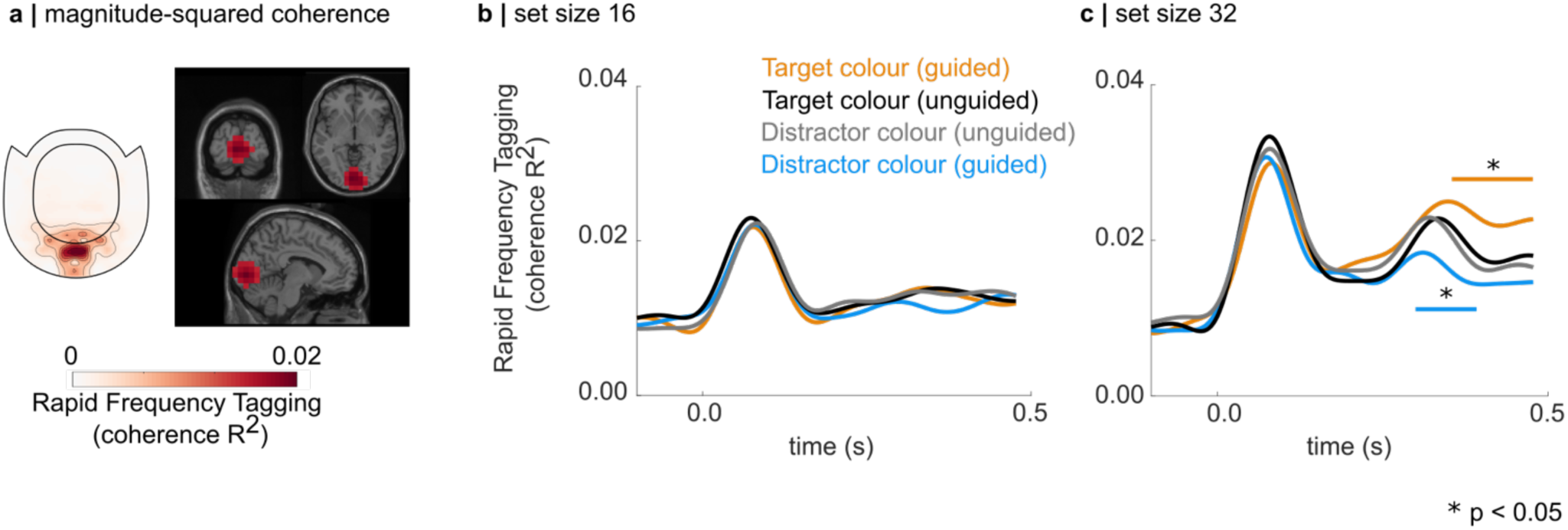
Rapid Invisible Frequency Tagging (RIFT) responses reflect a priority-map based mechanism. **a** (left) Topographic representation of the 60 Hz RIFT signal, averaged over participants in the 0 to 0.5 s interval (t = 0 s is the onset of the search display). The RIFT response is confined to the occipital sensors. (right) Source modelling demonstrates that the RIFT response was primarily generated in the early visual cortex. The source grid has been masked to show the 1% most strongly activated grid points (MNI coordinates [0 -92 -4]). **b,c** Coherence to the RIFT signal reveals target boosting and distractor suppression in guided search. **b** set size 16. There was no difference in RIFT responses between target and distractor colours; nor guided versus unguided search. **c** set size 32. The RIFT responses to the guided target colour are significantly enhanced and the responses to the guided distractor colour are significantly reduced compared to the unguided search condition (p < 0.05; multiple comparison controlled using a cluster-based permutation test in the 0 to 0.5 s interval).

As outlined in our pre-registration (https://osf.io/vcshj), we hypothesised that the RIFT response reflects a priority-map-based search strategy, indicating target boosting and distractor suppression in the *guided search* condition. Figure 2b and c show the RIFT response quantified by the coherence (R^2^) between the MEG response (*RIFT sensors of interest*) and the frequency tagging signal, averaged over participants (see Method for details on the RIFT analysis). Note that the immediate increase in coherence after the search display onset reflects a broad-band evoked response, rather than the frequency-specific flicker signal. The RIFT responses for *set size 16* were noticeably weak and did not show any modulation to the target and distractor colour (Figure 2b). Considering that the coherence drops to baseline at ∼200 ms, we argue that this is the result of an insufficient signal-to-noise ratio caused by a comparably small number of pixels flickering.

For *set size 32*, we find that the RIFT responses to the target colour in the *guided search* condition were significantly enhanced compared to the *unguided search* condition (average of the black and grey lines in Figure 2c, p < 0.05; multiple comparisons were controlled using a Monto-Carlo cluster-based dependent sample t-test on the 0 to 0.5 interval, 1,000 permutations). This suggests a boosting of the neuronal excitability to all items sharing the known target colour. Importantly, the responses to the distractor colour when comparing *guided* to *unguided search* were significantly reduced, providing evidence for distractor suppression (Figure 2c, p < 0.05; 1,000 permutations). Our findings demonstrate that knowledge about the target and distractor colour in the *guided search* condition results in a modulation of the RIFT response consistent with the concept of a priority map reflected in early visual cortex, whereby target representations are boosted, and distractor representations are suppressed. Importantly, we show that RIFT is suitable to measure the neuronal excitability associated with the priority map in visual search (also see Bouwkamp et al., 2023).

We further tested whether this modulation was relevant for performance, by sorting the trials based on a median split of reaction time into fast and slow trials. Importantly, faster responses were associated with a larger difference between the target and distractor response early in the trial, implying that successful visual search relies on a rapid modulation of the neuronal excitability in line with the concept of a priority map. These analyses are described in detail in the Supplementary analyses and Supplementary Figure 2.

Even though participants were instructed to perform the search task without moving their eyes, an argument could be advanced that the observed modulation of the RIFT responses results from an eye movement bias towards the target. We accounted for this potential confound based on a median split on reaction time (see Supplementary analyses). As demonstrated in Supplementary Figure 3, fast trials were not associated with a significantly higher or lower number of blinks or saccades compared to slow trials. While we did ensure that the search stimuli of each colour were evenly randomised over the display, we further tested whether the participants tended to move their gaze towards the target colour. This gaze bias was identified by binning the single trial eye tracking data into 100ms intervals, and counting how often the gaze was closest to a stimulus in the target colour. The number of occurrences the gaze was closest to the target colour was then divided by the total number of bins in the trial and averaged over all trials. As shown in Supplementary Figure 3c, the gaze bias appeared to average about 0.5 (i.e. 50% of the time bins within a trial), indicating that the gaze was in the vicinity of target and distractor stimuli about equal amounts of time. Importantly, there was no difference in gaze bias for fast vs slow trials, suggesting that participants generally followed the instructions and solved the task without moving their eyes. We conclude that the target boosting and distractor suppression observed in the RIFT response reflects a modulation of the excitability of early visual neurons in line with a priority-map based mechanism, that underlies visual attention and not eye movement.

### Investigating target boosting and distractor suppression at the single-trial level using a Generalized Linear Model

The mean-squared coherence results described above quantify the degree to which the variance in the MEG signal is accounted for by the RIFT signal (Cohen, 2014; Pan et al., 2021). This makes it a more interpretable measure for RIFT than spectral power, which is likely to be confounded by changes in broadband and/or high-frequency oscillatory activity. One caveat of coherence, however, is that it requires averaging over time (C. G. Richter et al., 2015). A single-trial quantification of the RIFT response is desirable, as it opens opportunities for several different analytic approaches, for instance, a Generalized Linear Model approach, used to link changes in oscillatory activity to experimental manipulations and behaviour (Griffiths et al., 2021; Quinn et al., 2024). Here, we developed a single-trial analysis of the RIFT response to investigate whether the presented priority map effect can be replicated using a single trial approach. Furthermore, using this approach, we were able to account for effects of task duration, and thus neural adaptation to the repeated presentation of the search display (Gardner et al., 2005).

To quantify the RIFT response at the single trial level, we first time-shifted the RIFT signal based on the onset of the steady-state visually evoked response, which has been shown to improve the signal-to-noise ratio (Spaak et al., 2024). The single-trial MEG data were first bandpass filtered at 60 and 67 Hz ± 3 Hz, and then averaged over trials (Figure 3a). We then fitted a sine wave to the 60 and 67 Hz response by setting all values of the filtered MEG signal higher than 0 to 1, and all values below zero to - 1. The resulting rectangular function (pink line in Figure 3a, top left) was then low-pass filtered at 110 Hz to obtain the phase-shifted sine wave (green line). To quantify the single trial RIFT response, we calculated the Spearman correlation between the phase-shifted sine wave (from 0.25 to 0.5 seconds after display onset) with the single trial MEG data. The resulting topography is shown for the combined planar gradiometers in Figure 3a bottom, showing the strongest correlation in a small set of occipital sensors.

**Figure 3.**
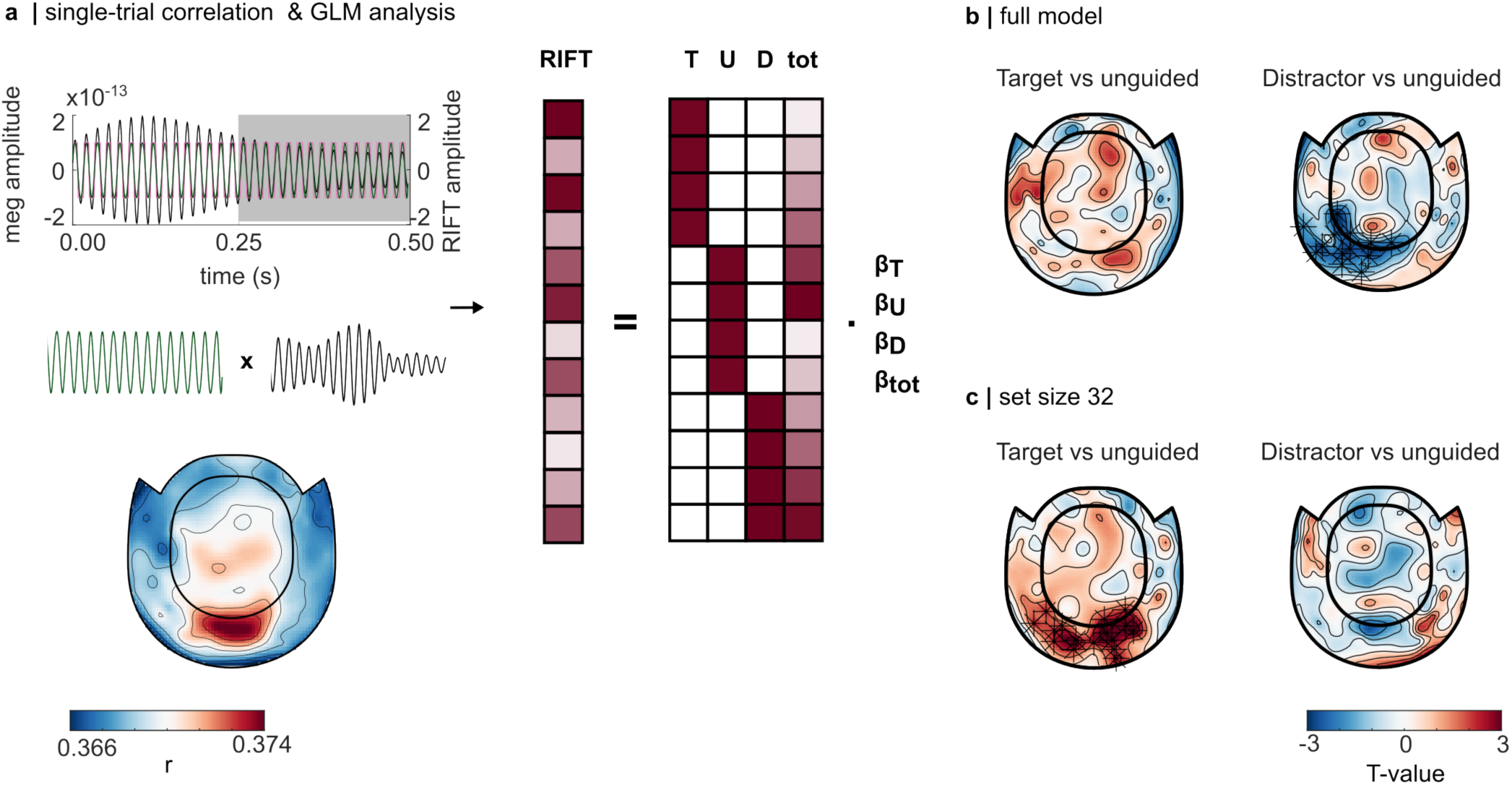
Quantifying the RIFT response at the single-trial level. **a** (top) To improve the signal-to-noise-ratio, we phase-shifted the RIFT signal based on the band-pass filtered steady-state response at each frequency (here shown for 60 Hz, black line). A perfect sine wave was fitted to the filtered signal by first setting all values above 0 to 1 and values below 0 to -1 (pink line). The edges were then removed by applying a low-pass filter (110 Hz) to the rectangular line. The single-trial RIFT response was quantified by correlating the single-trial MEG signal in each sensor (band-pass filtered at 55 to 75 Hz with the sine wave (left, bottom). The RIFT response to the target and distractor were then concatenated into one vector, and submitted to a GLM with the factors target colour (T), unguided (U), distractor colour (D), and time-on-task (tot). (bottom) The single-trial correlation in the combined planar gradiometers, indicating a strong RIFT response in the occipital sensors. **b** The contrast between the regressors associated with the target colour and unguided, and distractor colour and unguided, were compared to 0 using a cluster-based permutation test (5,000 permutations). The full model, including both set sizes, demonstrates significantly reduced responses to the distractor, while a model fit on set size 32 demonstrates evidence for target boosting.

For the GLM approach, the guided Target, guided Distractor, and unguided stimuli were modelled as separate regressors βT, βD, and βU, respectively (^F^igure 3a, right). Additionally, time-on-task (*tot*) was integrated into the design matrix based on the trial index for the concatenated RIFT responses for the targets and the distractors, and ranged from 0 to 1. To account for the inter-individual variability in the RIFT signal (Duecker et al., 2021), we fit the model to each participant individually, and calculated the T-values for target boosting and distractor suppression based on the contrast between βT and βU and βD and βU (see ^M^ethods). The resulting T-values in each MEG sensor, and for each participant, were then compared to 0 using a dependent-sample cluster-based permutation t-test.

For the full model, including *set size 16* and *set size 32*, we found that the contrast between βD and βU indicates significantly weaker RIFT responses to the distractor colour (Figure 3b, right). The difference between the βT and βU was, however, not significant ^(^Figure 3b, left). When fitting the GLM only to trials with *set size 32*, we found that the T-values associated with the βT and βU contrast were significantly larger than 0, indicating that the RIFT responses to the target were significantly enhanced compared to the *unguided search* condition (p < 0.05). The contrast between βD and βU, however, was not significant (p > 0.05).

These findings are generally in line with the coherence results presented above. The sustained target boosting shown by the coherence approach in set size 32 (orange line in Figure 2c) was replicated by the GLM. The RIFT responses in the *unguided search* condition (indicated by the black and grey line in Figure 2c) decrease over the duration of the search, which in turn reduces the difference to the RIFT response to the guided distractor colour. This explains the absence of evidence for distractor suppression for the GLM approach in the *set size 32* condition.

Overall, the GLM approach paired with the phase-shifted RIFT signal replicated the results of the mean-squared coherence. As such, the presented method can be used as a complementary measure to link RIFT responses to behaviour or other neural measures of interest, without the need to collapse over trials.

## Discussion

We used Rapid Invisible Frequency Tagging (RIFT) in combination with MEG as a novel approach to probe neuronal excitability in visual cortex, to investigate feature-guided visual search. In the *guided search* condition, the target colour was cued, whereas in the *unguided search* condition, the target colour was unknown. As expected, search performance was reduced for higher set sizes and for *unguided* compared to *guided* search; the latter confirming that participants used the colour cue at the beginning of the block to guide their search. Importantly, the RIFT responses revealed in the *guided search* condition, *set size 32*, demonstrated an increase in neuronal excitability in early visual cortex associated with the target colour and a suppression associated with the distractor colour. These results demonstrate that signs of a priority-map based mechanism, operating in a colour-specific manner, are observed in early visual regions. As we will outline below, this mechanism is likely to underlie top-down control by the ventral stream.

Our work complements previous electrophysiological recordings in humans and non-human primates investigating visual search paradigms with smaller set sizes of up to six items (Donohue et al., 2018; Forschack et al., 2022; Hickey et al., 2009; Ipata et al., 2009; Klink et al., 2023; van Zoest et al., 2021), and selective attention to moving stimuli (Andersen et al., 2008; Andersen & Müller, 2010; Müller et al., 2006). To achieve our aim, we employed a complex visual search display with a large set size, in the tradition of psychophysical research (Treisman et al., 1980; Wolfe, 1994, 2021). Recent studies have suggested that the RIFT signal does not propagate beyond V1/V2 (Duecker et al., 2021; also see Schneider et al., 2023; Soula et al., 2023), which is consistent with our source modelling results. We therefore propose that our findings provide evidence for a retinotopically organised priority map supported by early visual regions. The modulation of the RIFT signal can be observed at about 200 ms after search display onset. This time course is congruent with the observation that guidance by colour takes about 200-300 ms to be effective (E. M. Palmer et al., 2019), and further conforms with the latency of previously observed effects of attention on neural activity in V1 (Roelfsema et al., 1998). Electrophysiological recordings in non-human primates have shown target boosting and distractor suppression in the frontal eye field and lateral intraparietal cortex about 90 ms after stimulus onset (Cosman et al., 2018; Ipata, Gee, Goldberg, et al., 2006; Ipata, Gee, Gottlieb, et al., 2006) and after about 110 ms in V4 (Klink et al., 2023). As such, we argue that the priority map may be implemented in higher-order areas, and modulate neuronal excitability in V1 through feedback connections (Chen & Seidemann, 2012; Hochstein & Ahissar, 2002; Kamiyama et al., 2016; Muckli, 2010; Muckli & Petro, 2013). Recruiting the early visual neurons in a complex search task may be beneficial as the small receptive fields allow a higher spatial resolution when guiding the search (Ahissar & Hochstein, 1997, 2004). To find the target in the presented search, participants further needed to distinguish between T’s and L’s, which are defined by the spatial arrangement of the vertical and horizontal bars. These visual stimulus properties have been argued to be processed in caudal-medial areas of the dorsal pathway (V7), which are strongly connected to early visual and ventral regions (Freud et al., 2016). As such, modulating the excitability in early visual areas may allow the visual system to efficiently integrate colour and spatial information of stimuli that are likely to be the target.

The time course of the modulated RIFT response further indicates that neural excitability to the distractor colour was not reduced in anticipation of the search display. While recent evidence suggests that expected distractor locations can be suppressed before the onset of the stimuli (D. Richter et al., 2024), the results presented here demonstrate that this might not be the case for a complex, colour-guided visual search task.

### Limitations & Outlook

We did not find evidence for target boosting and distractor suppression for the lower set size of 16 items. Considering that the coherence drops to baseline after the event-related, frequency-unspecific response, this most likely reflects an insufficient signal-to-noise ratio of the RIFT response, caused by a larger distance between the stimuli and half as many pixels flickering for *set size 16* compared to *32*. In future studies, it would be interesting to investigate to verify this assumption by placing the 16 stimuli on a smaller search array or increasing stimulus size.

Target boosting and distractor suppression have been argued to be implemented by two distinct mechanisms (Donohue et al., 2018, 2020; Noonan et al., 2016). In this study, participants were able to infer the distractor colour from the cue provided at the beginning of the block, thus preventing us from disentangling these mechanisms. In future studies, it would be useful to use RIFT in a paradigm relying solely on distractor inhibition (Thayer et al., 2022), in which the participants are only informed about the distractor colour, while the target colour varies. This would clarify how distractor suppression is implemented when the target colour is unknown.

## Conclusion

In conclusion, our work demonstrates that guided search is associated with a modulation of neuronal excitability in early visual regions according to a priority map. Based on previous work, we argue that this mechanism underlies top-down control from downstream region. We propose that guided search relies on a priority map affecting neuronal excitability as early as primary visual cortex.

## Methods

### Experimental design & stimuli

#### Task

We applied Rapid Invisible Frequency Tagging (RIFT) in a classic visual search paradigm to probe the neuronal excitability to the target and distractor colour in *guided* and *unguided search*. The participants’ task was to indicate whether a cyan or yellow letter “T” was present or absent amongst several cyan and yellow “Ls” (Figure 1a). The experiment was designed in blocks of 40 trials with set sizes of either 16 or 32 items. At the beginning of a block in the *guided search* condition, the letter “T” was presented in yellow or cyan, indicating the colour of the target for the following block (Figure 1a). In blocks in the *unguided search* condition a white “T” was shown before the trial, and the colour of the target in the search display, if present, was randomly chosen to be cyan or yellow over trials. Each search display was preceded by a 1.5-s baseline interval in which a white fixation dot was presented in the centre of the screen. The trials were terminated with the participants’ button press, or automatically after 4 seconds. The button press was followed by a black screen, presented for 500 ms, before the start of the pre-search interval of the following trial. All participants completed four practice blocks consisting of 10 trials each before the experiment. Participants were instructed to find the target without moving their eyes. The experiment and MEG recording were paused every 10 minutes and participants were encouraged to rest their eyes and move their heads.

### Display Physics

The stimuli were presented using a Propixx lite projector (VPixx Technologies Inc, Quebec, Canada), set to a refresh rate of 480 Hz. The luminance of the yellow and cyan stimuli in the search display was modulated sinusoidally, respectively at 60 and 67 Hz (Figure 1b, target and distractor colours, tagging frequencies, and set sizes were randomized within participants). The stimuli were created using the Psychophysics Toolbox version 3 (Brainard, 1997) in MATLAB 2017a (The Mathworks, Natick, MA, USA).

### Apparatus for data acquisition

The MEG data were acquired using a MEGIN Triux (MEGIN Oy, Espoo, Finland), with 204 planar gradiometers and 102 magnetometers at 102 sensor positions, housed in a magnetically shielded room (Vacuumschmelze GmbH & Co, Hanau, Germany). Data were filtered online between 0.1 and 330 Hz using anti-aliasing filters and then sampled at 1,000 Hz. The dewar orientation was set to 60° to allow the participants to comfortably rest their heads against the back of the sensor helmet, optimizing the recording of the neuromagnetic signals in the occipital cortex.

The three fiducial landmarks (nasion and left and right periauricular points), the participant’s head shape (>200 samples), and the location of four head-position-indicator (HPI) coils were digitized using a Polhemus Fastrack (Polhemus Inc, Vermont, USA) prior to the recording. The location of the HPI coils was acquired at the beginning of each new recording block, but not continuously throughout the experiment.

The RIFT signals at 60 and 67 Hz were further applied to two squares at the outer corners of the screen and recorded using two custom-made photodiodes (Aalto NeuroImaging Centre, Aalto University, Finland), connected to the MEG system.

Eye movements and blinks were tracked using an EyeLink® eye tracker (SR Research Ltd, Ottawa, Canada), positioned at the minimum possible distance from the participant. The conversion of the EyeLink® Edf files was done with the Edf2Mat Matlab Toolbox designed and developed by Adrian Etter and Marc Biedermann at the University of Zurich.

The T1-weighted anatomical scans were obtained using a whole-body 3-Tesla Philips Achieva scanner (echo time TE=0.002s, repetition time TR=2s).

### Participants

This study was carried out in accordance with the declaration of Helsinki and the COVID-19 related safety measures at the University of Birmingham in place between April 2021 and January 2022. A telephone screening was conducted 48 hours before the experiment to ensure that all participants were safe for MRI and free of COVID-19 symptoms. 48 volunteers with no history of neurological disorders participated in the experiment. The participants’ colour vision was assessed prior to the experiment using 14 Ishihara plates (Clark, 1924). Participants for whom the eye tracking recording was missing due to technical errors were not considered for the analysis (N=6). Three additional participants were excluded as their button presses often extended into the following trials (in 160-300 trials), resulting in a total sample size of N=39. Participants who did not show a significant tagging response (N=8) were excluded at a later stage (see RIFT response sensor selection below), leaving 31 data sets (20 female, see below)

### Behavioural performance

The participants’ performance on correctly detecting the presence and absence of the target was quantified based on average reaction time and perceptual sensitivity (d’), calculated as:

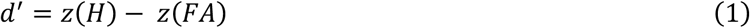

with z(H) being the z-scored portion of hits in target present trials and z(FA) being the z-scored portion of false alarms in target absent trials.

### MEG pre-processing

Signal Space Separation (SSS, “Maxfilter”) implemented in MNE Python was applied to suppress magnetic signals emerging from sources outside the participant’s brain. The remaining pre-processing of the MEG data, frequency and source analyses, and cluster-based permutation test were performed using the Fieldtrip toolbox (Oostenveld et al., 2010) in MATLAB 2019b. Statistical analyses of the behavioural and eye tracking data were carried out in RStudio 1.1.456 with R version 3.6.1. (The R Foundation for Statistical Computing).

Faulty sensors were identified and corrected prior to the SSS using MNE python. The filtered data were divided into intervals of 4.5 s, starting 2.5 s before, and extending to 2s after the onset of the search display in each trial. Semi-automatic artefact rejection was performed on the 4.5 s intervals, by manually identifying and rejecting epochs with a comparably high variance, separately for gradiometers and magnetometers. Independent Component Analysis (ICA) was used to suppress oculomotor and cardiac artefacts based on the 68 components that were identified for each participant. Trials with unreasonably short reaction times of up to 200ms, as well as trials without a response were rejected (Wolfe et al., 2010).

### RIFT response magnitude

#### Mean-squared coherence

For the offline analyses, we replaced the photodiode signals with a perfect sine wave with low-amplitude white noise (SE= 0.05) for the offline analyses, extending into the baseline interval. The magnitude of the RIFT response was quantified by calculating the spectral coherence between the MEG sensors of interest, identified as described above, and RIFT signal. The data were bandpass-filtered using a two-pass Butterworth filter at 60 and 67 Hz ± 5 Hz, respectively. The analytic signal was obtained from the filtered data using the Hilbert transform. The spectral coherence was then calculated as (Cohen, 2014, pp. 343– 344):

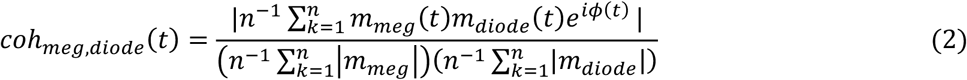

with m_meg_ and m_diode_ being the analytic MEG and RIFT amplitude, respectively, φ being the phase difference between the two signals, and n being the number of trials. To obtain the coherence to the RIFT signal of the target colour, for instance, we split the data into trials in which the target colour was tagged at 60 and 67 Hz, and calculated the coherence separately over these trials. Afterwards, the coherence was averaged over the two frequencies.

### RIFT response sensor selection

The MEG sensors containing a reliable frequency tagging response were identified using nonparametric (Monte Carlo) statistical testing, proposed by Maris & Oostenveld, 2007) and implemented in the Fieldtrip toolbox. The pre-processed data were divided into a baseline (0.7 to 0.2 s before stimulus onset) and stimulation interval (0.5 s following the onset of the search display). Coherence between a given MEG sensor and the 60 Hz photodiode signal over trials was estimated separately for the pre-search and the search interval. The difference between the coherence in the baseline and search interval was z-transformed using the following equation:

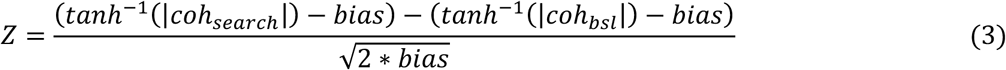

Whereby coh_search_ and coh_bsl_ are the coherence between the respective MEG sensor and the photodiode at 60 Hz during the search and pre-search interval, respectively. The bias is calculated as 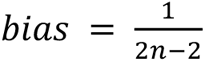 with n being the number of trials.

The statistical significance of the z-transformed coherence difference (the empirical z-value) was estimated using a permutation procedure. To this end, a null distribution for the empirical z-value was estimated by generating 10,000 random permutations of the trial labels and calculating the z-values for the shuffled pre-search and search interval, again using equation (3). If the coherence difference obtained for the unshuffled data in the respective sensor was larger than 99% of the null distribution, the sensor was considered to show a significant tagging response at a 1% significance level. This procedure was completed for a total of 81 occipital and occipito-parietal sensors to identify the sensors of interest for each participant. 31 out of 39 participants had at least one significant gradiometer. As only 27 participants showed a significant response in at least one magnetometer, only gradiometers were considered for the sensor and source analyses. In total, we used the data from 31 volunteers for further analyses (20 female; aged 23.4 years ± 3.18). All participants were right-handed according to the Edinburgh Inventory (augmented handedness score: M=84.08; STD=14.37, ref (Oldfield, 1971)).

### Generalized Linear Model spectrum

As the mean-squared coherence averages over observations (see Equation (2)), we next sought to develop a single-trial quantification of the RIFT response. To enhance the signal-to-noise ratio, we first calculated a phase-shifted measure of the RIFT signal (Spaak et al., 2024). The single-trial MEG signal was band-pass filtered at 60 and 67 Hz ± 3 Hz over the -2 to 4 second interval to avoid filter ringing. A sinewave was then fitted to the band-pass filtered steady state response by first fitting a rectangular function to the signal, setting values below 0 to -1 and values above 0 to 1, which was then low-pass filtered at 110 Hz to remove the edges (Figure 3a). To get the single-trial RIFT measure, we band-pass filtered the single-trial MEG data at 55 to 75 Hz, and calculated the Spearman correlation between the phase-shifted RIFT signal and MEG signal from 0.25 to 0.5 s after stimulus onset.

The single trial RIFT measures for the targets and distractors were then concatenated for each participant and further investigated using a Generalized Linear Model (GLM) approach (for an in-depth description of the method see Quinn et al., 2024).

The single-trial correlation in each gradiometer (*RIFT_grad_*) was modelled as a linear function of stimulus identity according to

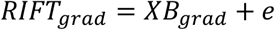

*RIFT_grad_* contained the RIFT responses in the gradiometers, concatenated for targets and distractors, respectively. *X* is the design matrix with four columns for the guided target, guided distractor, and unguided stimuli, respectively, as well as time-on-task. The target column contained values of 1 for each row with a RIFT response to a guided target, and 0 for the unguided stimuli and distractors. Column 2 and 3 followed the same logic and were set to 1 for the unguided stimuli and distractors, respectively, and 0 everywhere else. Time-on-task values ranged from 0 to 1.

*RIFT_grad_* contains the regressors associated with the target colour, unguided, and distractors for each gradiometer and was estimated by multiplying the Moore-Penrose pseudo-inverse of *X* with *RIFT_grad_* (Quinn et al., 2024)

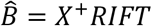

note that we dropped the *grad* index for readability.

Following conventional approaches in fMRI data analysis (Friston et al., 1994), we calculated the Contrast of Parameter Estimates (*cope*) for the GLM are calculated by multiplying the estimated regressors with the contrast vector *C*

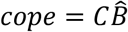

whereby 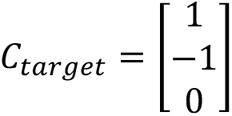 was multiplied with B̑ to quantify target boosting and 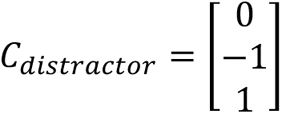 was used to quantify distractor suppression. The variance of the respective contrast of the model was calculated as

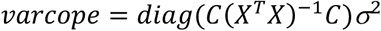

With σ^2^ being the variance of the residuals. The t values for each contrast were then calculated as

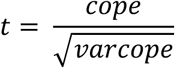

We used the t values rather than the estimated regressors, as they account for the variance of the contrasts (Quinn et al., 2024). The resulting t-values for each sensor were then compared to 0 using a one-tailed cluster-based permutation t-test with 5,000 permutations.

### Source localization

The anatomical sources of the RFT response were estimated using the Dynamic Imaging of Coherent Sources (Gross et al., 2001) beamformer, implemented in the Fieldtrip toolbox (Oostenveld et al., 2010)

### MEG lead field

To calculate the MEG lead field, we first aligned the fiducial landmarks in the individual T1-weighted images with the digitized points taken prior to the experiment. The coordinate system of the participant’s T1-weighted scan was then automatically aligned to the digitized head shape using the iterative closest point (ICP) algorithm(Besl & McKay, 1992), implemented in the Fieldtrip toolbox, and corrected manually as necessary. For the two participants for whom there was no T1 scan available, the digitized fiducial landmark and head shape were aligned with a standardized template brain provided with the Fieldtrip toolbox.

Next, the brain volume was discretized into a source grid of the equivalent current dipoles by warping each participant’s realigned anatomical scan to the Montreal Neurologic Institute (MNI) coordinate system; using a template MRI scan, and an equally spaced 8 mm grid, with 5,798 locations inside the brain. The lead field was then estimated at each point in the source grid using a semi-realistic headmodel(Nolte, 2003).

### Dynamic Imaging of Coherent Sources (DICS)

The spatial filters of the DICS beamformer were calculated as a function of the forward model (estimated using the lead field matrix) and the cross-spectral density matrix of the sensor data. Here, we used the cross-spectral matrix of the gradiometers only. The SSS (“Maxfilter”) caused the data to be rank deficient, making the estimate of the sensor cross-spectral density matrix unreliable. To ensure numerical stability, we calculated the truncated singular value decomposition (SVD) pseudoinverse (Gencer & Williamson, 1998; Westner et al., 2022) of the sensor cross-spectral density matrix. This method decomposes the covariance matrix using SVD, selects a subset of singular values (the subset size is defined by the numerical rank) and calculates a normalized cross-spectral density matrix using this subset. The spatial filters are then estimated based on the normalized cross-spectral density matrix using unit-noise gain minimum variance beamforming (Borgiotti & Kaplan, 1979; Westner et al., 2022).

To estimate the cross-spectral density matrix for the RIFT response, we first extracted data segments from 0 to 0.5 s (the minimum reaction time for all participants). The complex cross-spectral density between the signal in the (uncombined) planar gradiometers and the RIFT signal was computed based on the Fourier-transformed data segments (Hanning taper, separately for the 60 Hz and the 67 Hz photodiode signal). The cross-spectral density matrices were used to estimate the forward model to create a spatial filter for each frequency. The spatial filters were then applied to the cross-spectral density matrix to estimate the RIFT response as the coherence between each point in the source grid and the photodiode signal.

The spatial filters of the alpha oscillations were estimated based on the cross spectral density matrix of the gradiometers at the individual alpha frequency, calculated based on the -1 to 0 s interval (Hanning taper). Analogously to the RIFT response, the filters were than applied to calculate power at the individual alpha frequency at each grid point.

## Supplementary Material

**Supplementary Figure 1.**
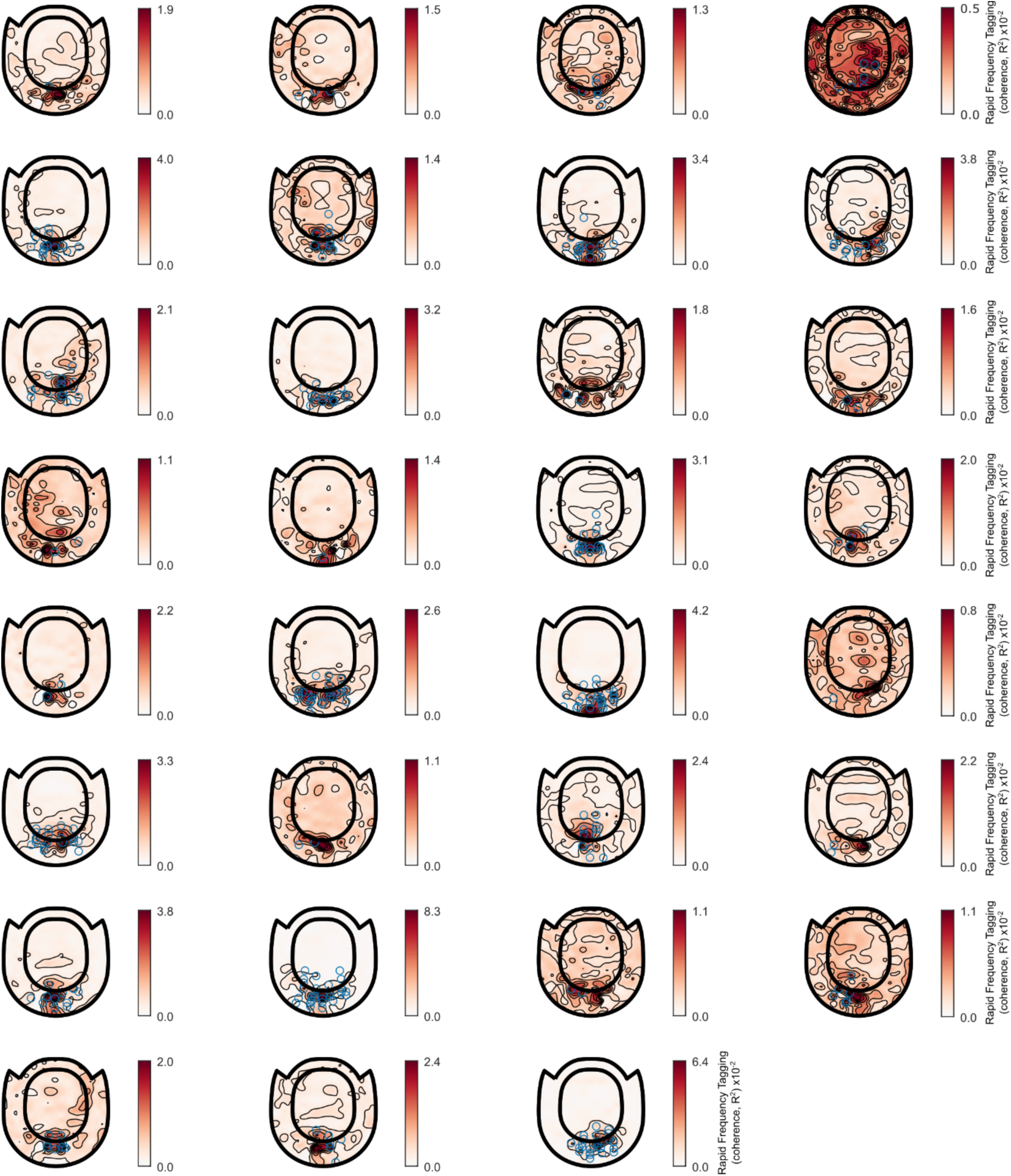
Topoplots showing coherence to RIFT at 60 Hz for each participant. RIFT sensors of interest are indicated by the light blue rings.

**Supplementary Figure 2.**
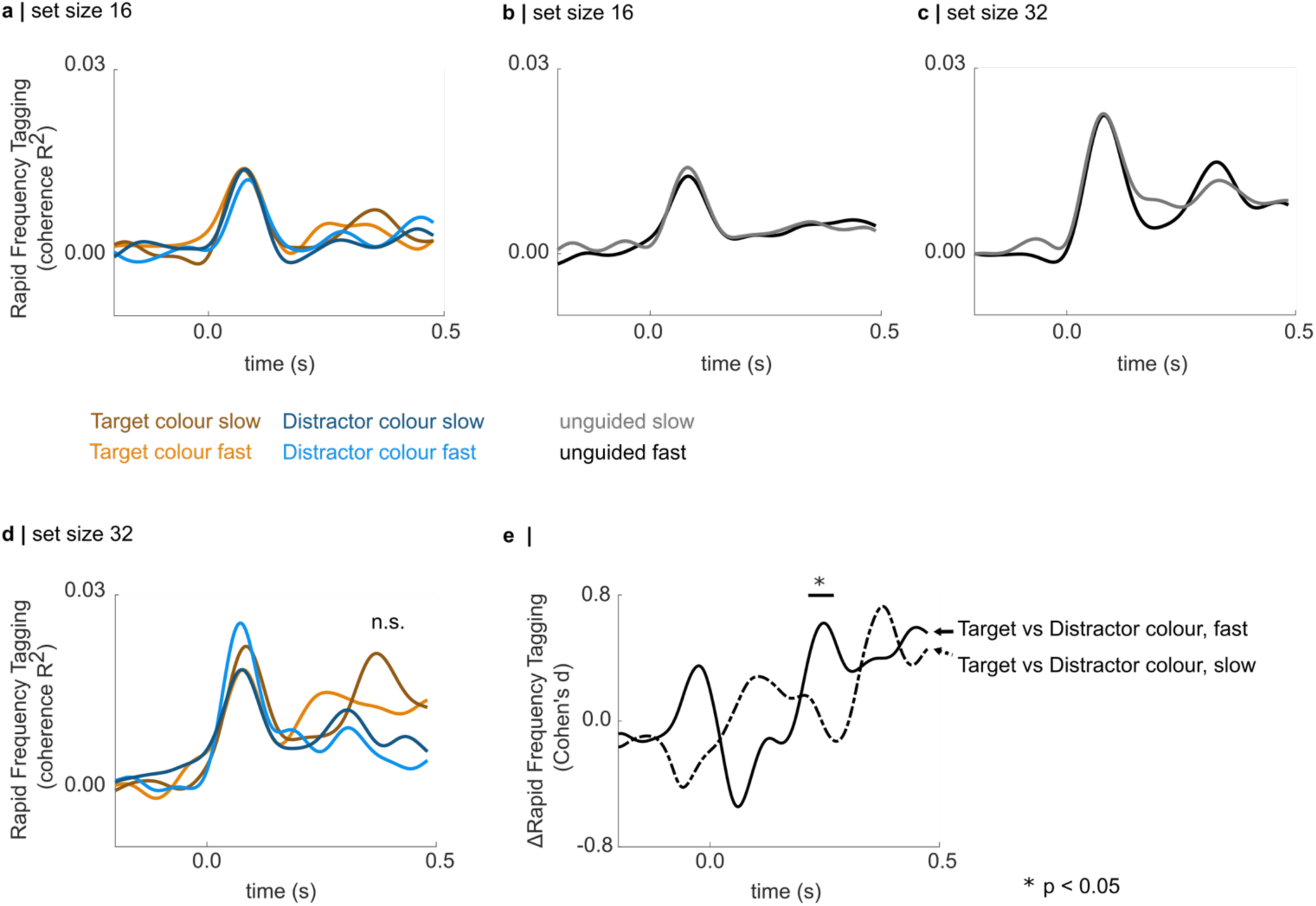
RIFT responses for fast and slow trials. There is no modulation of the RIFT response for fast vs slow trials for guided search, set size 16 (**a**) or unguided search set size 16 (**b**) or 32 (**c**). **d** Visual inspection of the RIFT responses in guided search set size 32 suggests higher coherence for fast compared to slow trials, however, neither the comparisons between targets for fast vs slow nor for the distractors revealed a significant difference (cluster-based permutation t-test, 1,000 permutations). **e** Upon further inspection of the interaction effects, we find that fast trials are associated with a larger difference in the RIFT response between the target and distractor colour (cluster-based permutation Monte Carlo t-test, p < 0.05, 1,000 permutations). This effect seems to mainly be driven by an initial increase in RIFT response to the target colour.

**Supplementary Figure 3.**
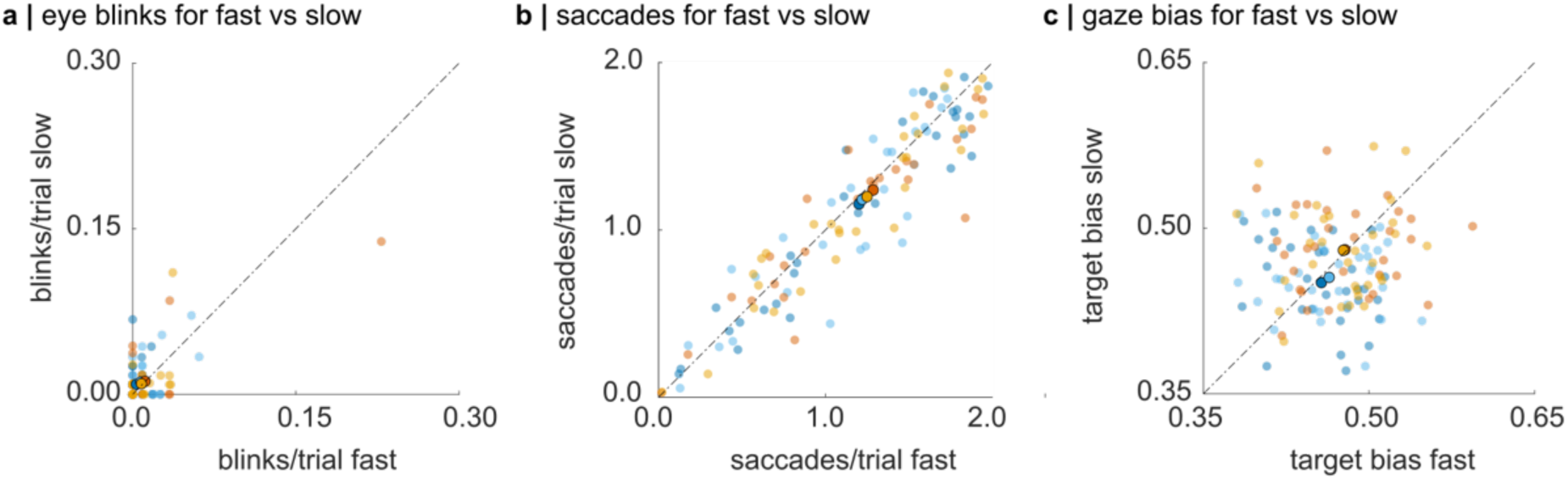
Eye blinks, saccades, and gaze bias towards the target colour for fast vs slow trials. Individual participants are indicated by the opaque scatters, the grandaverage is shown by the solid scatters. **a** Average number of eye blinks in each condition compared for fast and slow trials. A hierarchical regression approach revealed that the factor fast/slow did not add significant explanatory value to the baseline model including subject-specific random effects (χ^2^(1) = 1.9, p = 0.17). **b** Average number of saccades per trial during the search interval in each condition for fast vs slow trials. A hierarchical regression approach indicates a significant main effect for fast vs slow compared to the baseline model (χ^2^(1) = 3.9, p = 0.047, ΔR^2^ = 0.0014), however, the pairwise comparisons did not show any significant differences (Supplementary Table 5) **c** Proportion of the trial the gaze spent near the target colour. A hierarchical regression approach did not show a significant main effect for fast/slow, confirming that participants’ gaze tended to spend an equal amount of time within each trial close to stimuli in the target and distractor colour, as indicated by the value of 0.5 (χ^2^(1) = 0.19, p = 0.665).

**Supplementary Table 1.** Hierarchical regression on reaction time, revealing a significant main effect for set size and guided/unguided, but no interaction effect. When fitting the regression models, we assumed that the reaction time data followed a gamma distribution.

| Model | AIC | $\chi^2$ | p | R<sup>2</sup> |
| --- | --- | --- | --- | --- |
| $RT = \beta_0 + \varepsilon_{subj}$<br>$\varepsilon_{subj}$ subject-specific random effect | -117.04 | | | 0.59 |
| $RT = \beta_0 + \beta_1 X_{set\_size} + \varepsilon_{subj\_setsize}$<br>with $X_{set\_size} = \begin{cases} 1, & \text{set size 16} \\ 2, & \text{set size 32} \end{cases}$<br>$\varepsilon_{subj\_setsize}$ subject-specific, set size related random effect | -181.72 | 70.68 | $3 \times 10^{-15}$ | 0.77 |
| $RT = \beta_0 + \beta_1 X_{set\_size} + \beta_2 X_{guided} + \varepsilon_{subj\_setsize} + \varepsilon_{subj\_guided}$<br>with $X_{guided} = \begin{cases} 0, & \text{unguided} \\ 1, & \text{guided} \end{cases}$<br>$\varepsilon_{subj\_guided}$ subject-specific, guided/unguided related random effect | -307.57 | 133.86 | $2 \times 10^{-16}$ | 0.96 |
| $RT = \beta_0 + \beta_1 X_{set\_size} + \beta_2 X_{guided} + \beta_3 X_{interaction} + \varepsilon_{subj\_setsize} + \varepsilon_{subj\_guided}$ | -306.08 | 0.51 | 0.47 | 0.96 |

**Supplementary Table 2.** Reaction time contrasts between conditions (dependent sample Wilcoxon signed rank test). SE indicates the standard of the difference between means. Bonferroni corrected, * p < 0.05, ** p < 0.01, *** p < 0.001, **** p < 0.0001

|  | **unguided**<br>**search,**<br>**set size 16** | **guided search,**<br>**set size 16** | **unguided**<br>**search,**<br>**set size 32** |
| --- | --- | --- | --- |
| **guided search,**<br>**set size 16** | $V(30) = 481$<br>$z = 4.57$<br>$p = 8 \times 10^{-6}****$<br>$r = 0.82$<br>$SE_{diff} = 0.020$ | | |
| **unguided**<br>**search,**<br>**set size 32** | $V(30) = 5$<br>$z = 4.76$<br>$p = 6 \times 10^{-7}****$<br>$r = 0.86$<br>$SE_{diff} = 0.020$ | $V(30) = 0$<br>$z = 4.86$<br>$p = 1 \times 10^{-11}****$<br>$r = 0.87$<br>$SE_{diff} = 0.03$ | |
| **guided search,**<br>**set size 32** | $V(30) = 134$<br>$z = 2.34$<br>$p = 0.148 \text{ n.s.}$<br>$r = 0.4$<br>$SE_{diff} = 0.019$ | $V(30) = 0$<br>$z = 4.86$<br>$p = 1 \times 10^{-11}****$<br>$r = 0.87$<br>$SE_{diff} = 0.017$ | $V(30) = 473$<br>$z = 4.41$<br>$p = 4 \times 10^{-5}****$<br>$r = 0.79$<br>$SE_{diff} = 0.02$ |

**Supplementary Table 3.** Hierarchical regression on accuracy (d’), revealing a significant main effect for set size and guided/unguided, but no interaction effect.

| Model | AIC | $\chi^2$ | p | R<sup>2</sup> |
| --- | --- | --- | --- | --- |
| $d' = \beta_0 + \varepsilon_{subj}$<br>$\varepsilon_{subj}$ subject-specific random effect | 307.10 | | | 0.48 |
| $d' = \beta_0 + \beta_{set\_size}X_{set\_size} + \varepsilon_{subj\_setsize}$<br>with $X_{set\_size} = \begin{cases} 1, set\ size\ 16 \\ 2, set\ size\ 32 \end{cases}$ | 261.56 | 47.54 | $5 \times 10^{-12}$<br>**** | 0.68 |
| $d' = \beta_0 + \beta_1X_{set\_size} + \beta_2X_{guided} + \varepsilon_{subj\_setsize}$<br>$+ \varepsilon_{subj\_guided}$<br>with $X_{guided} = \begin{cases} 0, unguided \\ 1, guided \end{cases}$ | 217.96 | 45.60 | $1 \times 10^{-11}$<br>**** | 0.8 |
| $d' = \beta_0 + \beta_1X_{set\_size} + \beta_2X_{guided} + \beta_3X_{interaction}$<br>$+ \varepsilon_{subj\_setsize} + \varepsilon_{subj\_guided}$ | 219.05 | 0.91 | 0.34 | 0.8 |

**Supplementary Table 4.**
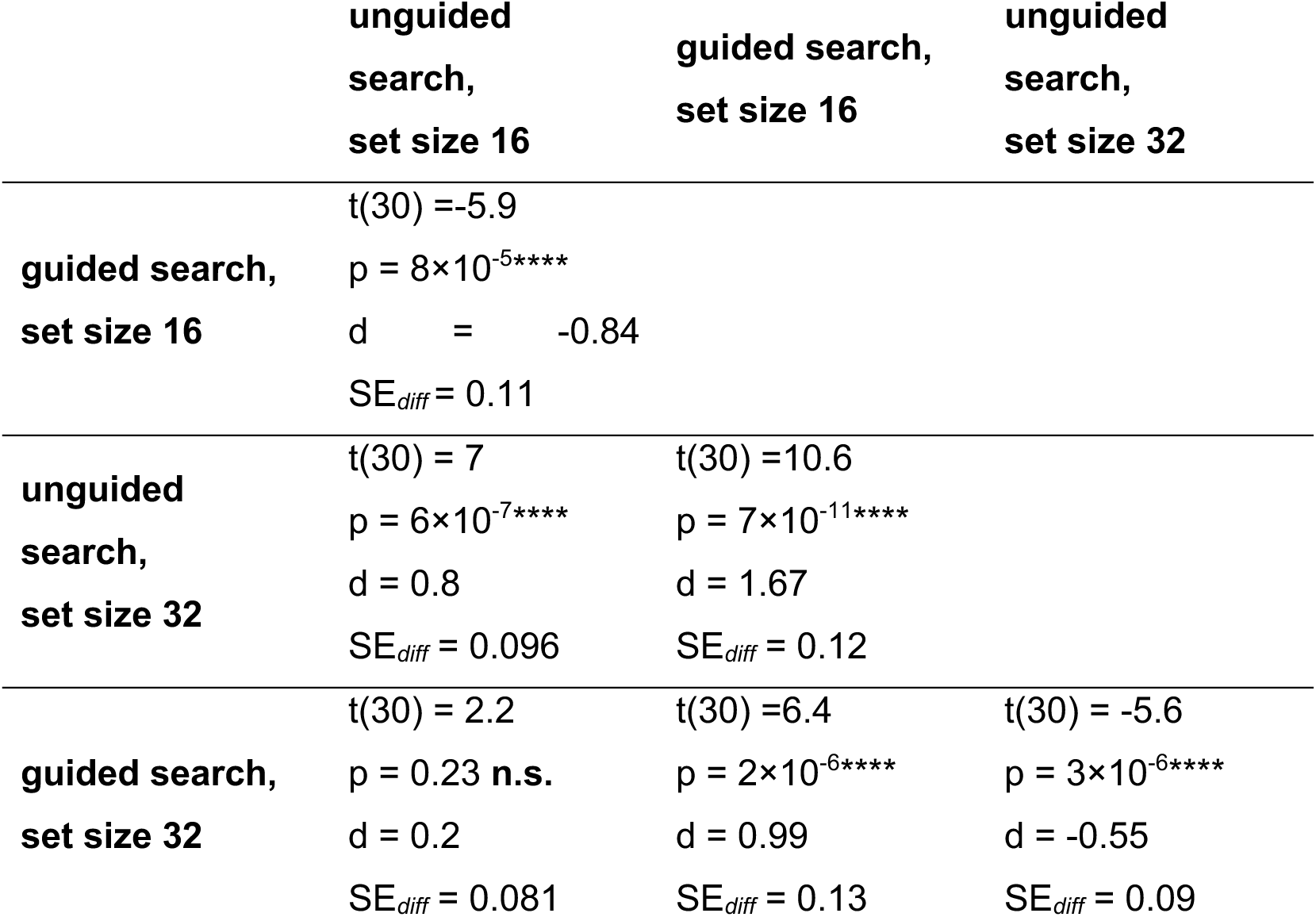
Sensitivity contrasts between conditions (post-hoc dependent sample t-tests, two-sided, Bonferroni-corrected).

**Supplementary Table 5.** Dependent sample t-tests on the number of saccades for fast vs slow trials reveals no significant effects of reaction time on saccades. The p-values are Benjamini-Hochberg corrected, however, none of the uncorrected p-values reached significance either.

| unguided search,<br>set size 16 | guided search,<br>set size 16 | unguided search,<br>set size 32 | guided search,<br>set size 32 |
| --- | --- | --- | --- |
| $t(30) = -0.92$<br>$p = 0.365$ | $t(30) = -1.7$<br>$p = 0.304$ | $t(30) = -1.47$<br>$p = 0.304$ | $t(30) = -1.21$<br>$p = 0.313$ |

## Supplementary analyses

### Behavioural results: Guided Search is associated with better performance

We predicted that search performance would become worse for more difficult search displays. Indeed, Figure 1c and d, suggests a decrease in reaction time, and an increase in accuracy (d’) for *guided* compared to *unguided search* and smaller (16) compared to larger (32) set sizes. We investigated these effects using a hierarchical regression approach with linear mixed models, whereby we consecutively added the factors *set size* and *guided/unguided* (dummy coded as 1 being “guided search”) into a model including only subject-related random effects.

To account for the skewed distribution of the reaction time data, we used a gamma distribution to fit the linear mixed models. A model predicting reaction time using the fixed factors *set size* and *guided/unguided*, and the random effects associated with these factors in each participant (AIC = -307.57), was superior to a model predicting reaction time as a function of subject-related random effects and *set size* (AIC = -181.72, χ^2^(4) = 133.86, p < 0.0001, R^2^ = 0.96, ΔR^2^ = 0.22). This additive model reveals a fixed effect for *set size* (β = 0.180) and *guided/unguided* (β = - 0.138). In summary, this demonstrates an increase in reaction time for set size 32 to compared to 16 by about 180ms, and an increase of about 138ms for un*guided* compared to *guided search* (no interaction effect, see Supplementary Table 1).

Analogously, hierarchical regression of accuracy shows that d’ is best predicted by an additive model including the factors *set size* (β = - 0.74) and *guided/unguided* (β = 0.56, R^2^ = 0.83) and the subject-specific random effects (see Supplementary Table 3). These results show that accuracy decreases for larger set sizes and increases for *guided* compared to *unguided search*.

The results of all post hoc tests are shown in Supplementary Table 2 and Supplementary Table 4, and indicated in Figure 1c and d. Notably, performance was not significantly different for *unguided search*, set size 16 and *guided search*, *set size 32* (Wilcoxon signed rank test on reaction time : V(30)=134, z = 2.23, p=0.148, r = 0.4, SE = 0.019; sensitivity (d’), dependent sample t-test: t(30)=2.2, p=0.23, d = 0.2, SE = 0.081). This finding is in line with the notion that *guided search* allows the participants to focus their search on items in the target colour, while ignoring the distractor colour.

In summary, the behavioural findings are consistent with a priori expectations, namely an increase in response times with set size, as well as faster responses for *guided* compared to *unguided search*.

### RIFT responses for fast compared to slow trials

As the analyses of the RIFT responses in *guided* and *unguided search* revealed a modulation of neuronal excitability in line with the priority map, we asked if successful target boosting and distractor suppression were relevant for performance. We therefore sorted the trials in each condition according to fast and slow responses (median split on reaction time) and compared the respective RIFT signals. Supplementary Figure 2d shows the RIFT response to the target colour for fast and slow trials (orange and brown line, respectively), and to the distractor colour (light and dark blue for fast and slow trials, respectively) for *guided search*, *set size 32*. We expected fast trials to be associated with respectively a stronger response to the target colour and a weaker response to the distractor colour, however, we did not find any significant differences between the RIFT responses to targets and distractors for fast vs slow trials. Upon further post-hoc comparisons, using cluster-based Monte Carlo permutations, we did find that the difference between the RIFT responses to the target and distractor colour is significantly larger for fast compared to slow trials, at about 200 ms after search display onset (Supplementary Figure 2e, p < 0.05, 1000 permutations). We conclude that there is a relationship between reaction time and the RIFT responses to target and distractor features, that is mainly driven by a strong initial response to the target colour.

### Ocular artefacts and gaze bias do not relate to RIFT responses

While participants were instructed to perform the task without moving their eyes, we found that some eye movements were present during the search. As enhanced neural processing has been suggested to underlie microsaccades (Liu et al., 2022; Lowet et al., 2018), we investigated if the reaction time effects and the modulated RIFT responses observed for fast vs slow trials can be explained by differences in ocular artefacts (Supplementary Figure 3).

We divided the trials in the eye tracking data in each condition based on the median reaction time, separately for target present and absent trials in each participant (as described in the main text). Next, we identified the number of blinks and saccades in the first 500 ms after the search display onset (the time interval included in the RIFT analyses) and averaged these over conditions. Note that the threshold of the eye tracker to identify a saccade was set to 0.6°. We again analysed all main and interaction effects using a hierarchical regression approach, by comparing the explanatory value of a model containing the factor fast/slow to the baseline model.

For the average number of blinks during the trial, we find that a model containing the factor *fast/slow* (AIC = -1259.3) did not explain a significantly larger portion of the variance than the baseline model including subject-specific random effects (AIC = -1259.5, χ^2^(1) = 1.9, p = 0.17 Supplementary Figure 3a). Using the same approach on the average number of saccades revealed that a model including the predictor *fast/slow* (AIC = -6.8) could indeed account for a larger portion of the variance than the baseline model (AIC = -4.86, χ^2^(1) = 3.9, p = 0.047, ΔR^2^ = 0.0014), however, none of the pairwise comparisons reached significance (Supplementary Table 5, Supplementary Figure 3b).

To ensure that any eye movements during the search were not over-proportionally directed at the target colour in the fast trials, we binned the eye tracking data into 100ms intervals and identified the stimulus closest to the location of the gaze in each of these bins. The gaze bias towards the target colour was defined as the proportion of time the eyes were directed at a position closest to a stimulus in the target colour. A value of 0.5 indicates that the participant’s gaze time on the target and the distractor colour were the same, meaning no bias. Comparing a linear regression model predicting gaze bias as a function of the factor *fast/slow* (AIC = -8.64.09), with a baseline model (AIC = - 865.9), did not reveal any significant main effects of reaction time on the gaze bias (χ^2^(1) = 0.19, p = 0.665, ΔR^2^ = 0.0001). This reveals that participants followed the instructions and did not solve the task by moving their eyes towards the target colour (Supplementary Figure 3c).

## Author Contributions

S.H. proposed research idea, K.D., K.L.S., S.H., J.W., and O.J. designed the experiment, K.D. acquired and analysed the data, Y.P. and O.J. supported analysis of MEG data, B.J.G. supervised the implementation of the GLM approach, K.D. and O.J. wrote the paper, all authors edited the paper.

### Acknowledgements

The authors thank Veikko Jousmaki for providing the light-to-voltage converter, Jonathan L. Winter for support with the MEG data acquisition, and Katarzyna Dudzikowska, Brandon Ingram, Alexander Murray, Davide Aloi, and Nina Salman for performing the MRI scans.

## Notes

### Competing Interest Statement

The authors have declared no competing interest.

